## Supplemental Figures and Derivations for "Fixational Eye Movements Enhance the Precision of Visual Information Transmitted by the Primate Retina"

### Supplementary Information

#### Supplementary Figures

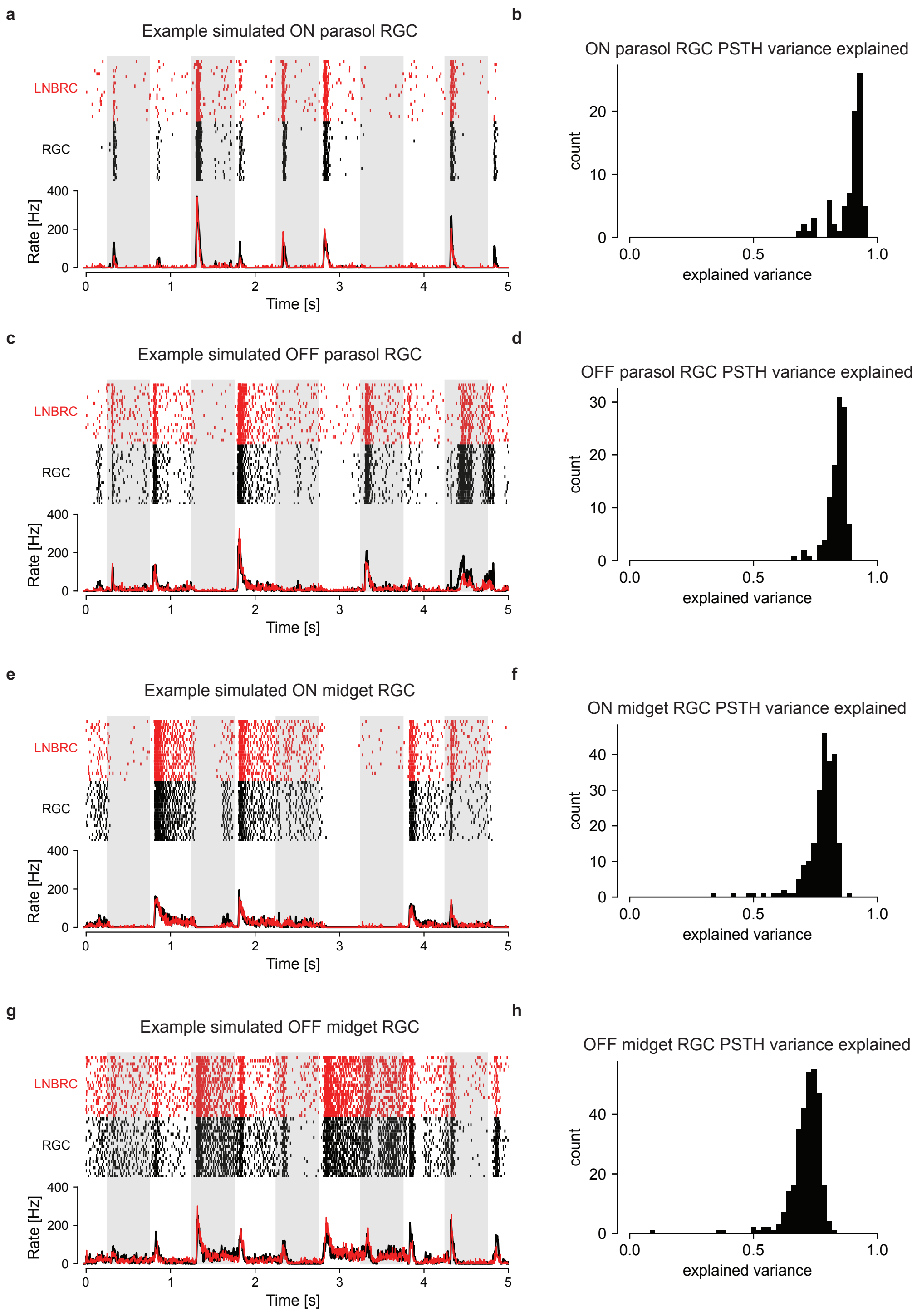


**Figure S1:** LNBRC fit quality for fixational drift natural movies, for cells of each of the major cell types in the macaque monkey (ON parasol, OFF parasol, ON midget, OFF midget). **(a)** Comparison of LNBRC-simulated repeat rasters and PSTH with real recorded repeats in response to fixational drift natural movie stimuli, for a single example ON parasol RGC. **(b)** Histogram showing fraction of PSTH variance explained by the LNBRCs, for every ON parasol RGC recorded in one preparation. The fraction of explained variance was systematically high for every cell, suggesting that LNBRCs can accurately represent retinal responses to natural movie stimuli. **(c)** and **(d)** Same as (a) and (b) for OFF parasol RGCs. **(e)** and **(f)** Same as (a) and (b) for ON midget RGCs. **(g)** and **(h)** Same as (a) and (b) for OFF midget RGCs.

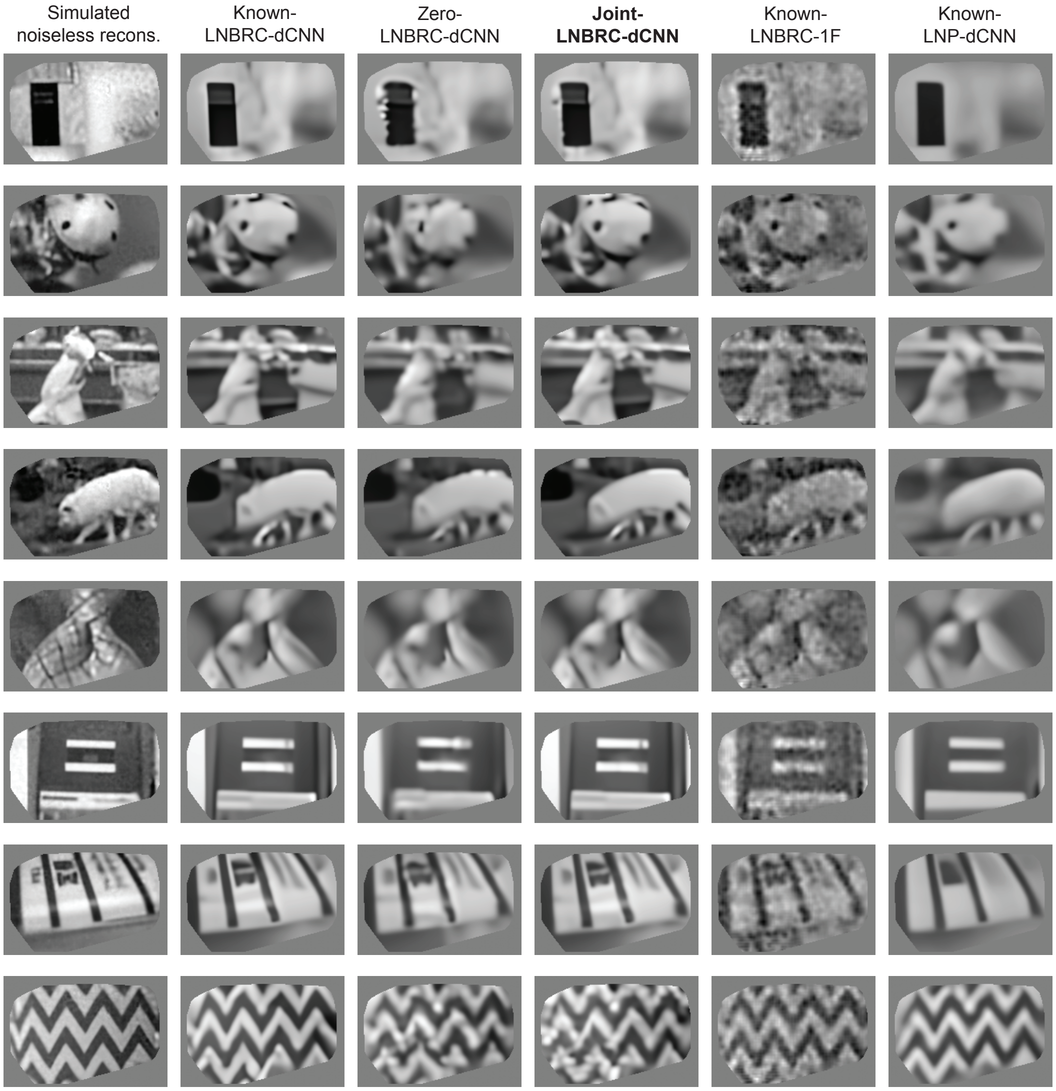


**Figure S2:** Additional example fixational drift stimulus images and reconstructions, from the same experimental preparation as Figure 2d from the main text, with additional comparisons against simpler reconstruction methods. Columns (left to right): Simulated noiseless reconstruction, a reconstruction of the stimulus from linear projections onto the LNBRC filters (see Methods); Known-LNBRC-dCNN, MAP reconstruction with known eye movements, using LNBRC encoding and dCNN prior; Zero-LNBRC-dCNN, MAP reconstruction while incorrectly assuming no eye movements, using LNBRC encoding and dCNN prior; Joint-LNBRC-dCNN, reconstruction by jointly estimating image and eye movements, using LNBRC encoding and dCNN prior; Known-LNBRC-1F, MAP reconstruction with known eye movements, using LNBRC encoding and simpler 1/F Gaussian prior; Known-LNP-dCNN, MAP reconstruction with known eye movements, using simpler LNP encoding and dCNN prior. The original stimuli were drawn from the ImageNet database [1], and are unavailable due to copyright restrictions.

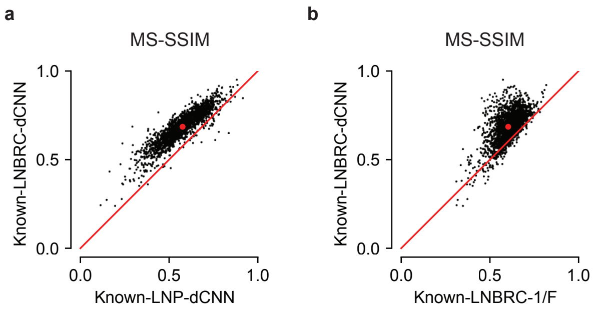


**Figure S3:** Performance comparison of the LNBRC-dCNN MAP reconstruction algorithm against simpler alternatives, for the eye movements natural movies, in the case that the eye movements are known *a priori*. $N = 1992$ test images were used to perform the comparisons in each panel. **(a)** Comparing LNBRC-dCNN against LNP-dCNN, in which the LNBRC encoding model was replaced with a simpler LNP encoding model. Reconstructions using the LNBRC encoding model were systematically better than those using the LNP model, demonstrating that the more sophisticated LNBRC encoding model contributed substantially to reconstruction quality. **(b)** Comparing LNBRC-dCNN against LNBRC-1/F, in which the dCNN prior was replaced with a simpler Gaussian 1/F prior. Reconstructions using the dCNN prior were systematically better than those using the 1/F prior, demonstrating that the dCNN natural image prior contributed substantially to reconstruction quality.

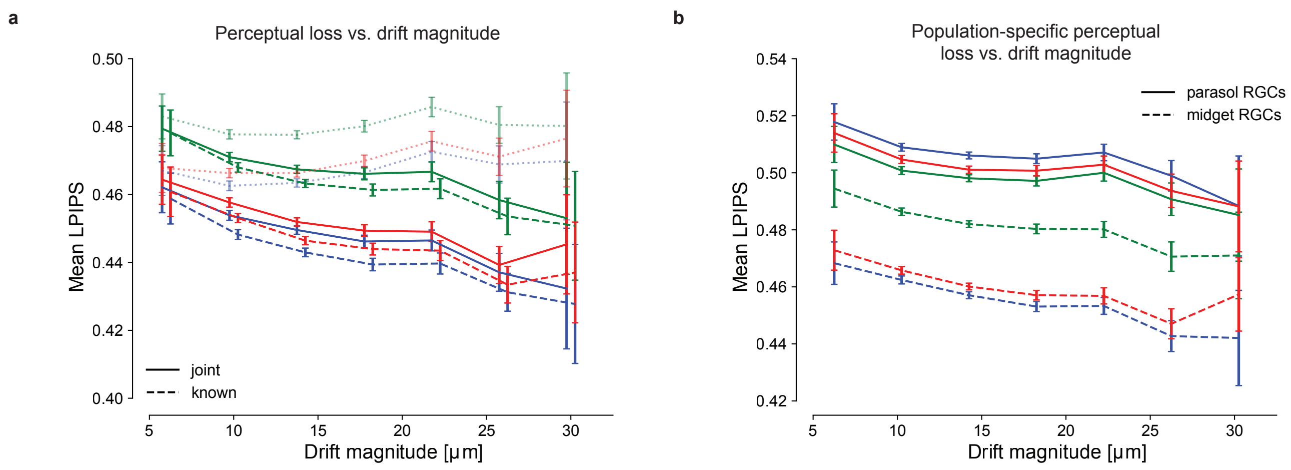


**Figure S4:** **(a)** Re-analysis of Figure 3a (reconstruction quality as a function of quantity of fixational drift eye movements) using LPIPS, an alternative measure of perceptual distance based on a deep neural network trained for object recognition. Smaller LPIPS values corresponds to higher quality. Each color corresponds to a different preparation (same color convention as Figure 3a). Error bars correspond to the standard error of the sample mean. The results using LPIPS are consistent with those using MS-SSIM: regardless of whether eye movements were known *a priori* (Known-LNBRC-dCNN, dashed lines) or jointly estimated along with the image (joint-LNBRC-dCNN, solid lines), LPIPS decreases with increasing eye movements (images improve in quality with increasing eye movements). Failure to compensate for eye movements (zero-LNBRC-dCNN, dotted lines) resulted in LPIPS increasing with increasing eye movements. **(b)** Re-analysis of Figure 3b (population-specific reconstruction quality as a function of quantity of fixational drift eye movements) using LPIPS. The solid lines correspond to the parasol-only joint-LNBRC-dCNN reconstructions, and the dashed lines to midget-only joint-LNBRC-dCNN reconstructions. Error bars correspond to the standard deviation of the sample mean. In all cases, LPIPS decreased with increasing eye movements, demonstrating that drift eye movements improved both the parasol cell and midget cell signals.

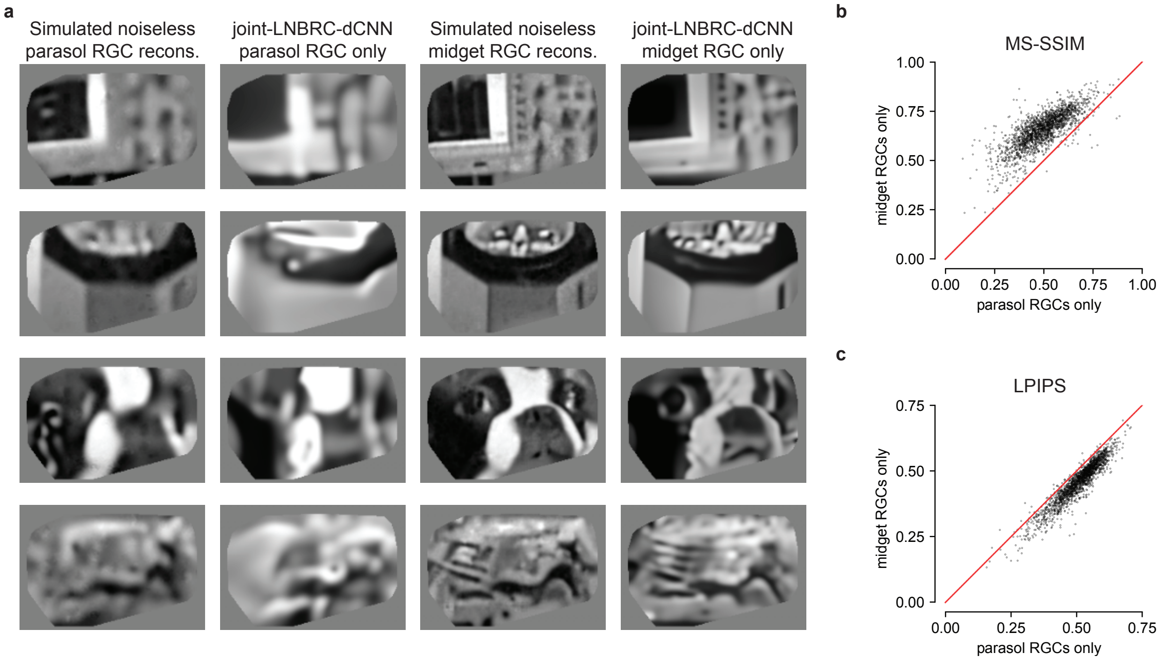


**Figure S5:** Comparison of parasol-only and midget-only reconstructions for the fixational drift eye movements stimulus, in one preparation. **(a)** Example reconstructions, using the joint-LNBRC-dCNN simultaneous eye movement estimation and image reconstruction algorithm. Columns (left to right): Simulated noiseless parasol RGC reconstruction, reconstructions of the stimuli from linear projections onto the parasol RGC LNBRC filters only (see Methods); joint-LNBRC-dCNN parasol RGC only, joint reconstructions computed from experimental data using only parasol RGCs; Simulated noiseless midget RGC reconstruction, reconstructions of the stimuli from linear projections onto the midget RGC LNBRC filters only (see Methods); and Midget RGCs, joint reconstruction from experimental data computed using only midget RGCs. The midget-only reconstructions contained greater fine spatial detail than the parasol-only reconstructions. The original stimuli were drawn from the ImageNet database [1], and are unavailable due to copyright restrictions. **(b)** Comparison of MS-SSIM reconstruction quality between the midget-only and parasol-only reconstructions. For nearly every image, the midget-only reconstruction quality exceeded that of parasol-only reconstructions. **(c)** Comparison of LPIPS perceptual distance between midget-only and parasol-only reconstructions. For nearly every image, the midget-only perceptual distance was smaller than that of parasol-only reconstructions.

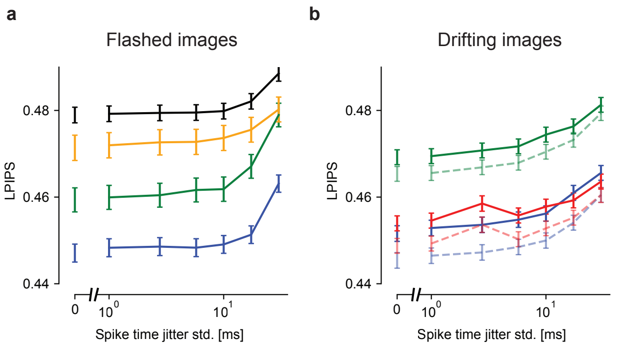


**Figure S6:** Re-analysis of Figure 4 using LPIPS. **(a)** Flashed natural image reconstruction performance as a function of spike timing perturbation, in four experimental preparations (colors). The x-axis is plotted on a log scale, with a broken axis to facilitate comparison with unperturbed data. Error bars in all panels correspond to the standard error of the sample mean. Performance at each level of temporal perturbation was evaluated using $N = 1500$, $N = 1750$, $N = 750$, and $N = 750$ images for the blue, black, green, and yellow preparations, respectively. LPIPS remained relatively constant up to 10 ms of spike timing jitter. **(b)** Fixational drift natural image reconstruction performance as a function of spike timing perturbation, in three experimental preparations. The solid lines correspond to joint reconstruction of the image and eye trajectory, while the dashed and faded lines correspond to reconstruction of the image alone with known eye trajectory. Performance at each level of temporal perturbation was evaluated using $N = 1992$ images for each experimental preparation. For the fixational drift stimulus, LPIPS gradually increased with increasing spike time perturbation, and degraded measurably with 2-5 ms of spike timing perturbation.

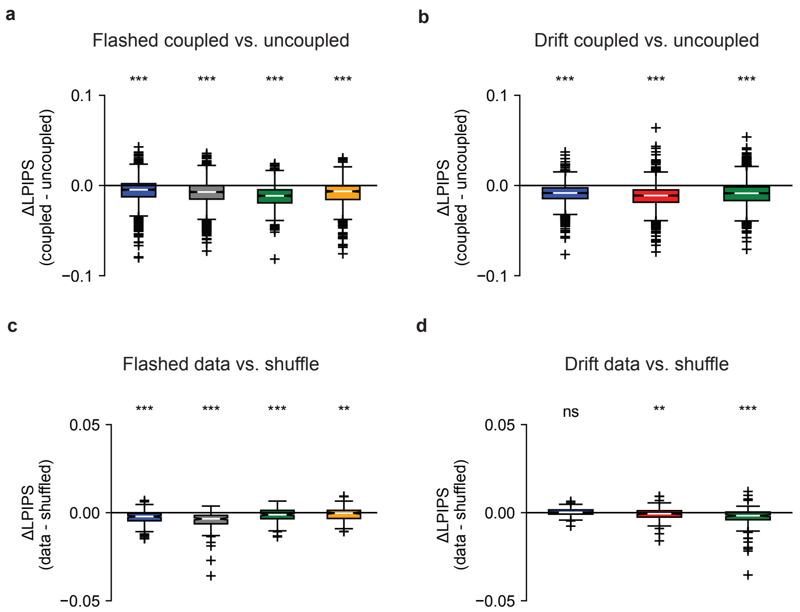


**Figure S7:** Re-analysis of Figure 5 using LPIPS. **(a)** For the flashed stimuli, LPIPS distances were significantly lower (better) when using the fully-coupled LNBRC models to perform reconstruction than when using the uncoupled LNBR models. From left to right, mean differences between coupled and uncoupled LPIPS were $-5.85 \cdot 10^{-3}$, $-8.20 \cdot 10^{-3}$, $-1.2 \cdot 10^{-2}$, $-8.54 \cdot 10^{-3}$ (p-values $1.7 \cdot 10^{-58}$, $6.7 \cdot 10^{-125}$, $1.1 \cdot 10^{-96}$, and $1.4 \cdot 10^{-54}$ respectively, Wilcoxon signed rank test, coupled < uncoupled, $N = 1500$, $N = 1750$, $N = 750$, and $N = 750$, respectively). For all boxplots (panels a-d), the box marks the median and the inter-quartile range (IQR), while the whiskers extend to 1.5 times the IQR. Outliers are marked with a +. **(b)** The same result held for the fixational drift stimulus. From left to right, mean differences between coupled and uncoupled LPIPS were $-8.69 \cdot 10^{-3}$, $-9.02 \cdot 10^{-3}$, and $-1.20 \cdot 10^{-2}$ (p-values $1.4 \cdot 10^{-210}$, $8.5 \cdot 10^{-169}$, and $1.3 \cdot 10^{-252}$, respectively, Wilcoxon signed rank test, coupled < uncoupled, $N = 1992$ for all). **(c)** For the flashed stimuli, LPIPS decreased when RGC responses were shuffled across repeats to remove noise correlations, but by a much smaller amount than when the coupling filters were removed. From left to right, mean LPIPS differences between data and shuffled reconstructions were $-2.73 \cdot 10^{-3}$, $-3.48 \cdot 10^{-3}$, $-1.16 \cdot 10^{-3}$, and $-1.85 \cdot 10^{-4}$ (p-values $5.4 \cdot 10^{-15}$, $6.8 \cdot 10^{-24}$, $5.45 \cdot 10^{-5}$, and $8.52 \cdot 10^{-3}$ respectively, coupled < uncoupled, Wilcoxon signed rank test, $N = 150$ for all). **(d)** The same result held for the fixational drift stimulus. From left to right, mean difference between data and shuffled reconstructions were $2.50 \cdot 10^{-4}$, $-7.06 \cdot 10^{-4}$, and $-2.55 \cdot 10^{-3}$ (p-values 0.946, $2.31 \cdot 10^{-3}$, and $6.14 \cdot 10^{-9}$ respectively, coupled < uncoupled, Wilcoxon signed rank test, $N = 149$ for all).

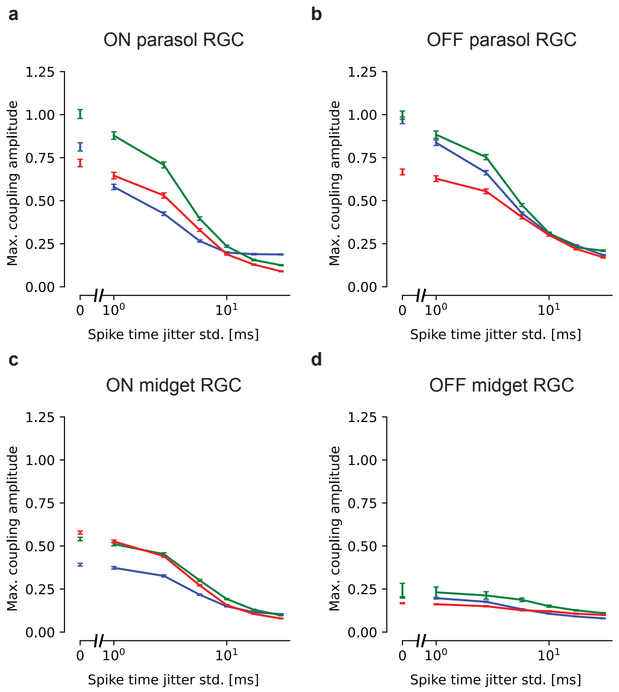


**Figure S8:** Mean peak amplitude of the LNBRC coupling filter for the nearest homotypic neighbor as a function of spike timing perturbation, in three preparations. The mean is taken across all RGCs of the specified type, and the error bars represent the standard error. **(a)** Peak amplitude of ON parasol RGC coupling filter to nearest neighbor ON parasol RGC. **(b)** Peak amplitude of OFF parasol RGC coupling filter to nearest neighbor OFF parasol RGC. **(c)** Peak amplitude of ON midget RGC coupling filter to nearest neighbor ON midget RGC. **(d)** Peak amplitude of OFF midget RGC coupling filter to nearest neighbor OFF midget RGC.

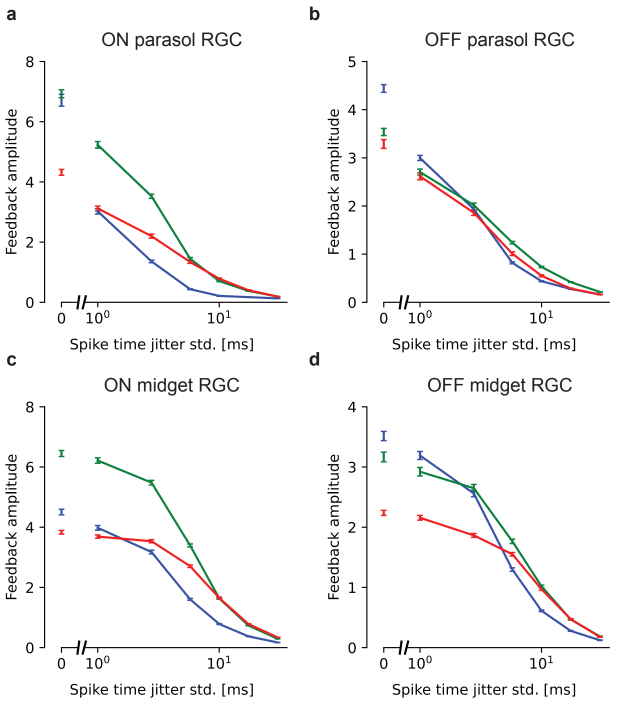


**Figure S9:** Mean peak amplitude of the LNBRC feedback filter as a function of spike timing perturbation, in three preparations. The mean is taken across all RGCs of the specified type, and the error bars represent the standard error. **(a)** Peak amplitude of ON parasol RGC feedback filter. **(b)** Peak amplitude of OFF parasol RGC feedback filter. **(c)** Peak amplitude of ON midget RGC feedback filter. **(d)** Peak amplitude of OFF midget RGC feedback filter.

### Supplementary Methods

#### 2.1 Derivations for MAP-EM reconstruction

Let $\mathbf{s}$ be the observed spike, $\mathbf{y}$ the static stimulus image, and $\mathbf{w}$ the full eye movement trajectory over all timesteps. In this case $\mathbf{w}$ is discrete (integer) valued rather than continuous, as the stimulus presented to the retina consisted of discretely shifted frames. The eye movements are assumed to satisfy the Markov property: letting $\mathbf{w}_t$ denote the eye position at timestep $t$, $p(\mathbf{w}_t \mid \mathbf{w}_0, \ldots, \mathbf{w}_{t-1}) = p(\mathbf{w}_t \mid \mathbf{w}_{t-1})$.

The encoding model is fitted to the data with full knowledge of the stimulus (e.g. with exactly known eye movements), and hence the encoding model is parameterized as $p(\mathbf{s} \mid \mathbf{y}, \mathbf{w})$. The exact MAP reconstruction problem is then formulated as

$$
\begin{aligned}
\arg\max_{\mathbf{y}} \big\{ \log p(\mathbf{y} \mid \mathbf{s}) \big\} &= \arg\max_{\mathbf{y}} \big\{ \log p(\mathbf{y}, \mathbf{s}) \big\} \\
&= \arg\max_{\mathbf{y}} \left\{ \log \sum_{\mathbf{w}_{0,\ldots,T}} p(\mathbf{y}, \mathbf{s}, \mathbf{w}) \right\} \\
&= \arg\max_{\mathbf{y}} \left\{ \log \sum_{\mathbf{w}_{0,\ldots,T}} p(\mathbf{s} \mid \mathbf{y}, \mathbf{w}) p(\mathbf{w}) p(\mathbf{y}) \right\} \\
&= \arg\max_{\mathbf{y}} \left\{ \log p(\mathbf{y}) + \log \sum_{\mathbf{w}_{0,\ldots,T}} p(\mathbf{s} \mid \mathbf{y}, \mathbf{w}) p(\mathbf{w}) \right\}.
\end{aligned}
$$

This optimization cannot be computed directly because marginalization over $\mathbf{w}$ (marginalization over all of the possible eye movement trajectories) is NP-hard and not tractable. We therefore introduce a variational distribution $q(\mathbf{w} \mid \mathbf{y}^{(i)}, \mathbf{s})$ on the eye movements, and use the EM algorithm to iteratively approximate the solution (see Supplementary Note 3.1 for derivation) with

$$
\hat{\mathbf{y}}^{(i+1)} = \arg\max_{\mathbf{y}} \big\{ \log p(\mathbf{y}) + \mathbb{E}_{\mathbf{w} \sim q(\mathbf{w} \mid \mathbf{y}^{(i)}, \mathbf{s})} \big[ \log p(\mathbf{s} \mid \mathbf{y}, \mathbf{w}) \big] \big\} \tag{2.1}
$$

This optimization problem can be solved using a Plug-and-Play algorithm, provided that the variational distribution $q$ is simple enough to easily compute the expectation over.

#### 2.2 Particle filter variational distribution $q$

The variational distribution $q(\mathbf{w} \mid \mathbf{s}, \mathbf{y})$ in the MAP-EM problem can be chosen arbitrarily, but should be close to $p(\mathbf{w} \mid \mathbf{s}, \mathbf{y})$ for the variational lower bound to be good. We choose define $q(\mathbf{w} \mid \mathbf{s}, \mathbf{y})$ to fit the form

$$q(\mathbf{w} \mid \mathbf{s}, \mathbf{y}) = \frac{p(\mathbf{s} \mid \mathbf{w}, \mathbf{y}) r_0(\mathbf{w}_0) \prod_{i=1}^{T} r_i(\mathbf{w}_i \mid \mathbf{w}_{i-1})}{p(\mathbf{s} \mid \mathbf{y})}$$

where $r_i(\mathbf{w}_i \mid \mathbf{w}_{i-1})$ can be an arbitrary distribution. This strongly resembles the form of the exact expression $p(\mathbf{w} \mid \mathbf{s}, \mathbf{y}) = \frac{p(\mathbf{s}|\mathbf{w},\mathbf{y})p(\mathbf{w}|\mathbf{y})}{p(\mathbf{s}|\mathbf{y})} = \frac{p(\mathbf{s}|\mathbf{w},\mathbf{y})p(\mathbf{w}_0)\prod_{i=1}^{T} p(\mathbf{w}_i|\mathbf{w}_{i-1})}{p(\mathbf{s}|\mathbf{y})}$.

Rather than build an expression for $q(\mathbf{w} \mid \mathbf{s}, \mathbf{y})$ all at once, we use a particle filter to represent $q^{(j)}(\mathbf{w} \mid \mathbf{s}, \mathbf{y})$, our estimate of $q$ after the first $j$ frame transitions, and iteratively update the particle filter once for each frame transition. $q^{(j)}$ must satisfy the functional form for $q$, and can be written as

$$q^{(j)}(\mathbf{w} \mid \mathbf{s}, \mathbf{y}) = \frac{p(\mathbf{s} \mid \mathbf{w}, \mathbf{y}) r_0^{(j)}(\mathbf{w}_0^{(j)}) \prod_{i=1}^{T} r_i^{(j)}(\mathbf{w}_i^{(j)} \mid \mathbf{w}_{i-1}^{(j)})}{p(\mathbf{s} \mid \mathbf{y})}$$

In order to update the particle filter for timestep $j+1$, we work with the unnormalized distribution (e.g. the numerator in the above expression) $\gamma_j = p(\mathbf{s} \mid \mathbf{w}, \mathbf{y}) r^{(j)}(\mathbf{w}_0^{(j)}) \prod_{i=1}^{T} r^{(j)}(\mathbf{w}_i^{(j)} \mid \mathbf{w}_{i-1}^{(j)})$. Using a sampling distribution $v_{j+1}(\mathbf{w}^{(j+1)} \mid \mathbf{w}^{(j)})$, the particle filter update weight $\alpha_{i+1}$ can be expressed as

$$\begin{aligned}
\alpha_{j+1} &= \frac{\gamma_{j+1}(\mathbf{w}_{0,\ldots,T}^{(j)})}{\gamma_j(\mathbf{w}_{0,\ldots,T}^{(j)}) v_{j+1}(\mathbf{w}^{(j+1)} \mid \mathbf{w}^{(j)})} \\
&= \frac{p(\mathbf{s} \mid \mathbf{w}^{(j+1)}, \mathbf{y}) r_0^{(j+1)}(\mathbf{w}_0^{(j+1)}) \prod_{i=1}^{T} r_i^{(j+1)}(\mathbf{w}_i^{(j+1)} \mid \mathbf{w}_{i-1}^{(j+1)})}{p(\mathbf{s} \mid \mathbf{w}^{(j)}, \mathbf{y}) v_{j+1}(\mathbf{w}^{(j+1)} \mid \mathbf{w}^{(j)}) r_0^{(j)}(\mathbf{w}_0^{(j)}) \prod_{i=1}^{T} r_i^{(j)}(\mathbf{w}_i^{(j)} \mid \mathbf{w}_{i-1}^{(j)})}
\end{aligned}$$

Note that we are yet to define what $r$ and $v$ are, and that we are free to define these however we wish. Let us define $v_{j+1}$ as

$$v_{j+1}(\mathbf{w}^{(j+1)} \mid \mathbf{w}^{(j)}) = \begin{cases} p(\mathbf{w}_{j+1} \mid \mathbf{w}_j = \mathbf{w}_j^{(j)}), & \mathbf{w}_{j+2,\ldots,T}^{(j+1)} = \mathbf{w}_{j+1}, \mathbf{w}_{0,\ldots,j} = \mathbf{w}_{0,\ldots,j}^{(j)} \\ 0 & \text{otherwise} \end{cases}$$

This definition means that we update the particle for timestep $j+1$ by setting the eye position at $\mathbf{w}_{j+1}$ as well as all subsequent eye positions to the same random draw from the distribution $p(\mathbf{w}_{j+1} \mid \mathbf{w}_j)$, while leaving all eye postions from previous times $t \leq j$ unchanged.

We then define $r_i^{(j)}(\mathbf{w}_i^{(j)} \mid \mathbf{w}_{i-1}^{(j)})$ to be the following:

- For $i = 0$ and any value of $j$,

$$r_0(\mathbf{w}_0^{(j)}) = \begin{cases} 1, & \mathbf{w}_0^{(j)} = \mathbf{0} \\ 0 & \text{otherwise} \end{cases}$$

- For $i \leq j$,

$$r_i^{(j)}(\mathbf{w}_i^{(j)} \mid \mathbf{w}_{i-1}^{(j)}) = p(\mathbf{w}_i \mid \mathbf{w}_{i-1} = \mathbf{w}_{i-1}^{(j)})$$

- For $i > j$,

$$r_i^{(j)}(\mathbf{w}_i^{(j)} \mid \mathbf{w}_{i-1}^{(j)}) = \begin{cases} 1, & \mathbf{w}_i^{(j)} = \mathbf{w}_{i-1}^{(j)} \\ 0 & \text{otherwise} \end{cases}$$

Substituting this set of definitions into the expression for $\alpha_{j+1}$, we find that

$$\begin{aligned}
\alpha_{j+1} &= \frac{p(\mathbf{s} \mid \mathbf{w}^{(j+1)}, \mathbf{y})}{p(\mathbf{s} \mid \mathbf{w}^{(j)}, \mathbf{y})} \cdot \frac{r_0^{(j+1)}(\mathbf{w}_0^{(j+1)}) \prod_{i=1}^{T} r_i^{(j+1)}(\mathbf{w}_i^{(j+1)} \mid \mathbf{w}_{i-1}^{(j+1)})}{v_{j+1}(\mathbf{w}^{(j+1)} \mid \mathbf{w}^{(j)}) r_0^{(j)}(\mathbf{w}_0^{(j)}) \prod_{i=1}^{T} r_i^{(j)}(\mathbf{w}_i^{(j)} \mid \mathbf{w}_{i-1}^{(j)})} \\
&= \frac{p(\mathbf{s} \mid \mathbf{w}^{(j+1)}, \mathbf{y})}{p(\mathbf{s} \mid \mathbf{w}^{(j)}, \mathbf{y})} \cdot \frac{p(\mathbf{w}_{j+1}^{(j+1)} \mid \mathbf{w}_j^{(j)}) \prod_{i=1}^{j} p(\mathbf{w}_i^{(j)} \mid \mathbf{w}_{i-1}^{(j)}) \prod_{i=j+2}^{T} r_i^{(j+1)}(\mathbf{w}_i^{(j+1)} \mid \mathbf{w}_{i-1}^{(j+1)})}{v_{j+1}(\mathbf{w}^{(j+1)} \mid \mathbf{w}^{(j)}) \prod_{i=1}^{j} p(\mathbf{w}_i^{(j)} \mid \mathbf{w}_{i-1}^{(j)}) \prod_{i=j+1}^{T} r_i^{(j)}(\mathbf{w}_i^{(j)} \mid \mathbf{w}_{i-1}^{(j)})} \\
&= \frac{p(\mathbf{s} \mid \mathbf{w}^{(j+1)}, \mathbf{y})}{p(\mathbf{s} \mid \mathbf{w}^{(j)}, \mathbf{y})} \cdot \frac{p(\mathbf{w}_{j+1}^{(j+1)} \mid \mathbf{w}_j^{(j)})}{v_{j+1}(\mathbf{w}^{(j+1)} \mid \mathbf{w}^{(j)})} \\
&= \frac{p(\mathbf{s} \mid \mathbf{w}^{(j+1)}, \mathbf{y})}{p(\mathbf{s} \mid \mathbf{w}^{(j)}, \mathbf{y})}
\end{aligned}$$

since both $\prod_{i=j+1}^{T} r_i^{(j)}(\mathbf{w}_i^{(j)} \mid \mathbf{w}_{i-1}^{(j)})$ and $\prod_{i=j+2}^{T} r_i^{(j+1)}(\mathbf{w}_i^{(j+1)} \mid \mathbf{w}_{i-1}^{(j+1)})$ both have value 1 in the only places that the respective distributions are supported. Hence $\alpha_{j+1}$ can be calculated by computing two encoding log-likelihood values, each using all of the spikes, and then subtracting.

The overall setup leads to a sequential importance resampling procedure for updating the particle filter at each timestep:

*At time* $t = 0$

- Assign $\mathbf{w}_0 = \mathbf{0}$, since we are free to assume that the eye position starts at an arbitrarily-defined origin.
- Create $N$ equally weighted particles, all located at the origin.

*At time* $t \geq 1$

- For each particle, update the positions for time $t, \ldots, T$ by drawing from $p(\mathbf{w}_t \mid \mathbf{w}_{t-1} = \mathbf{w}_{t-1}^{(t-1)})$ and setting all positions $\mathbf{w}_{t,\ldots,T}$ to that value.
- Compute weight updates $\alpha_t = \frac{p(\mathbf{s} \mid \mathbf{w}_{0,\ldots,T}^{(t)}, \mathbf{y})}{p(\mathbf{s} \mid \mathbf{w}_{0,\ldots,T}^{(t-1)}, \mathbf{y})}$.
- Resample to obtain $N$ new equally-weighted particles.

The particle filter MAP-EM algorithm is

**Algorithm 1** Particle filter MAP-EM for reconstruction with eye movements

Initialize $N$ particles at the origin, $\mathbf{w}_{0:T} = (0, 0)$, each with probability $\frac{1}{N}$ to represent the distribution $q^{(0)}(\mathbf{w} \mid \mathbf{x}, \mathbf{s})$.
$\mathbf{y}^{(0)} \leftarrow \arg\max_{\mathbf{y}} \left\{ \log p(\mathbf{y}) + \mathbb{E}_{\mathbf{w} \sim q^{(0)}(\mathbf{w} \mid \mathbf{y}^{(i)}, \mathbf{s})} \left[ \log p(\mathbf{s} \mid \mathbf{y}, \mathbf{w}) \right] \right\}$
**for** $k \in \{1, 2, \ldots, K\}$ **do**
    $q^{(k)} \leftarrow$ Update-Particles()
    $\mathbf{y}^{(k)} \leftarrow \arg\max_{\mathbf{y}} \left\{ \log p(\mathbf{y}) + \mathbb{E}_{\mathbf{w} \sim q^{(k)}(\mathbf{w} \mid \mathbf{y}^{(k-1)}, \mathbf{s})} \left[ \log p(\mathbf{s} \mid \mathbf{y}, \mathbf{w}) \right] \right\}$
    Resample-Particles()
**end for**

Note that although this algorithm builds the eye movements trajectory distribution forwards in time, it is *not* an online algorithm because it assumes that all of the spikes corresponding to the stimulus image presentation are available. Thus the algorithm integrates over the entire stimulus presentation.

### Supplementary Note

#### 3.1 MAP-EM reconstruction with unknown eye movements

The MAP objective in the case of unknown eye movements is

$$\begin{aligned}\arg\max_{\mathbf{y}}\big\{\log p(\mathbf{y}\mid\mathbf{s})\big\} &= \arg\max_{\mathbf{y}}\big\{\log p(\mathbf{y})+\log p(\mathbf{s}\mid\mathbf{y})\big\}\\ &= \arg\max_{\mathbf{y}}\big\{\log p(\mathbf{y})+\log\sum_{\mathbf{w}_{0,\ldots,T}} p(\mathbf{s},\mathbf{w}\mid\mathbf{y})\big\}\\ &= \arg\max_{\mathbf{y}}\left\{\log p(\mathbf{y})+\log\left[\sum_{\mathbf{w}_{0,\ldots,T}} p(\mathbf{s}\mid\mathbf{y},\mathbf{w})p(\mathbf{w})\right]\right\}\end{aligned}$$

which requires an intractable marginalization over the eye movements $\mathbf{w}$. Suppose on iteration $i$ of the algorithm, we have a guess for the image $\mathbf{y}^{(i)}$. We introduce a variational distribution $q(\mathbf{w}\mid\mathbf{y}^{(i)},\mathbf{s})$ and simplify the troublesome second term

$$\begin{aligned}\log p(\mathbf{s}\mid\mathbf{y}) &= \log\sum_{\mathbf{w}_{0,\ldots,T}} p(\mathbf{s},\mathbf{w}\mid\mathbf{y})\\ &= \log\left[\sum_{\mathbf{w}_{0,\ldots,T}} p(\mathbf{s},\mathbf{w}\mid\mathbf{y})\frac{q(\mathbf{w}\mid\mathbf{y}^{(i)},\mathbf{s})}{q(\mathbf{w}\mid\mathbf{y}^{(i)},\mathbf{s})}\right]\\ &= \log\mathbb{E}_{\mathbf{w}\sim q(\mathbf{w}\mid\mathbf{y}^{(i)},\mathbf{s})}\left[\frac{p(\mathbf{s},\mathbf{w}\mid\mathbf{y})}{q(\mathbf{w}\mid\mathbf{y}^{(i)},\mathbf{s})}\right]\\ &= \log\mathbb{E}_{\mathbf{w}\sim q(\mathbf{w}\mid\mathbf{y}^{(i)},\mathbf{s})}\left[\frac{p(\mathbf{s}\mid\mathbf{y},\mathbf{w})p(\mathbf{w})}{q(\mathbf{w}\mid\mathbf{y}^{(i)},\mathbf{s})}\right]\\ &\geq \mathbb{E}_{\mathbf{w}\sim q(\mathbf{w}\mid\mathbf{y}^{(i)},\mathbf{s})}\left[\log\frac{p(\mathbf{s}\mid\mathbf{y},\mathbf{w})p(\mathbf{w})}{q(\mathbf{w}\mid\mathbf{y}^{(i)},\mathbf{s})}\right]\\ &= \mathbb{E}_{\mathbf{w}\sim q(\mathbf{w}\mid\mathbf{y}^{(i)},\mathbf{s})}\big[\log p(\mathbf{s}\mid\mathbf{y},\mathbf{w})+\log p(\mathbf{w})\big]\\ &\quad-\mathbb{E}_{\mathbf{w}\sim q(\mathbf{w}\mid\mathbf{y}^{(i)},\mathbf{s})}\big[\log q(\mathbf{w}\mid\mathbf{y}^{(i)},\mathbf{s})\big]\end{aligned}$$

where the inequality is due to Jensen's inequality. Substituting this lower bound back into the original objective function leaves

$$\begin{aligned}\arg\max_{\mathbf{y}}\Big\{&\log p(\mathbf{y})+\mathbb{E}_{\mathbf{w}\sim q(\mathbf{w}\mid\mathbf{y}^{(i)},\mathbf{s})}\big[\log p(\mathbf{s}\mid\mathbf{y},\mathbf{w})+\log p(\mathbf{w})\big]\\ &-\mathbb{E}_{\mathbf{w}\sim q(\mathbf{w}\mid\mathbf{y}^{(i)},\mathbf{s})}\big[\log q(\mathbf{w}\mid\mathbf{y}^{(i)},\mathbf{s})\big]\Big\}\end{aligned}$$

Since $\mathbf{y}^{(i)}$ is a fixed known constant from the previous iteration of the algorithm, the term $\mathbb{E}_{\mathbf{w}\sim q(\mathbf{w}|\mathbf{y}^{(i)},\mathbf{s})}[\log q(\mathbf{w} \mid \mathbf{y}^{(i)}, \mathbf{s})]$ is simply a constant and can be discarded from the optimization. In addition, since the term $\mathbb{E}_{\mathbf{w}\sim q(\mathbf{w}|\mathbf{y}^{(i)},\mathbf{s})}[\log p(\mathbf{w})]$ doesn't depend on the optimization variable $\mathbf{y}$, we can discard it as well. This leaves the problem

$$\arg\max_{\mathbf{y}} \Big\{ \log p(\mathbf{y}) + \mathbb{E}_{\mathbf{w}\sim q(\mathbf{w}|\mathbf{y}^{(i)},\mathbf{s})}\big[\log p(\mathbf{s} \mid \mathbf{y}, \mathbf{w})\big] \Big\} \tag{3.1}$$

We have freedom to pick the distribution $q(\mathbf{w} \mid \mathbf{y}^{(i)}, \mathbf{s})$, but what distribution gives us the tightest bound? As it turns out, the gap between the inequality and the equality is exactly the KL-divergence between $q(\mathbf{w} \mid \mathbf{y}^{(i)}, \mathbf{s})$ and $p(\mathbf{w} \mid \mathbf{y}, \mathbf{s})$

$$\begin{aligned}
D_{KL}(q(\mathbf{w} \mid \mathbf{y}^{(i)}, \mathbf{s}) \parallel p(\mathbf{w} \mid \mathbf{y}, \mathbf{s})) &= \mathbb{E}_{\mathbf{w}\sim q(\mathbf{w}|\mathbf{y}^{(i)},\mathbf{s})}\left[\log \frac{q(\mathbf{w} \mid \mathbf{y}^{(i)}, \mathbf{s})}{p(\mathbf{w} \mid \mathbf{y}, \mathbf{s})}\right] \\
&= \mathbb{E}_{\mathbf{w}\sim q(\mathbf{w}|\mathbf{y}^{(i)},\mathbf{s})}[\log q(\mathbf{w} \mid \mathbf{y}^{(i)}, \mathbf{s})] \\
&\quad - \mathbb{E}_{\mathbf{w}\sim q(\mathbf{w}|\mathbf{y}^{(i)},\mathbf{s})}\left[\log \frac{p(\mathbf{w}, \mathbf{s} \mid \mathbf{y})}{p(\mathbf{s} \mid \mathbf{y})}\right] \\
&= \log p(\mathbf{s} \mid \mathbf{y}) - \mathbb{E}_{\mathbf{w}\sim q(\mathbf{w}|\mathbf{y}^{(i)},\mathbf{s})}\left[\log \frac{p(\mathbf{w}, \mathbf{s} \mid \mathbf{y})}{q(\mathbf{w} \mid \mathbf{y}^{(i)}, \mathbf{s})}\right]
\end{aligned}$$

and hence we want to choose a distribution $q(\mathbf{w} \mid \mathbf{y}, \mathbf{s})$ to be as close to $p(\mathbf{w} \mid \mathbf{y}, \mathbf{s})$ as possible to make the lower bound tight. Obviously, explicitly computing $p(\mathbf{w} \mid \mathbf{y}, \mathbf{s})$ will be rather difficult, and hence we iteratively build an approximation to it using particle filtering.

#### 3.2 Sequential importance sampling

Define $\gamma_n(\mathbf{w}_{0:n})$ as an unnormalized distribution. Let $p(\mathbf{w}_n \mid \mathbf{w}_{n-1})$ be the hidden state transition kernel, and let $p(\mathbf{s}_n \mid \mathbf{w}_n)$ be the observation model.

For sequential importance sampling, we pick a straightforward distribution $q_n(\mathbf{w}_{0:n}) = q(\mathbf{w}_0)\prod_{k=1}^{n} q_k(\mathbf{w}_k \mid \mathbf{w}_{k-1})$ to sample from, and then compute an unnormalized weight for the sample

$$\begin{aligned}
W_n(\mathbf{w}_{0:n}) &= \frac{\gamma_n(\mathbf{w}_{0:n})}{q_n(\mathbf{w}_{0:n})} \\
&= \frac{\gamma_{n-1}(\mathbf{w}_{0:n-1})}{q_{n-1}(\mathbf{w}_{0:n-1})} \frac{\gamma_n(\mathbf{w}_{0:n})}{\gamma_{n-1}(\mathbf{w}_{0:n-1}) q_n(\mathbf{w}_n \mid \mathbf{w}_{n-1})} \\
&= W_{n-1}(\mathbf{w}_{0:n-1}) \frac{\gamma_n(\mathbf{w}_{0:n})}{\gamma_{n-1}(\mathbf{w}_{0:n-1}) q_n(\mathbf{w}_n \mid \mathbf{w}_{n-1})} \\
&= W_{n-1}(\mathbf{w}_{0:n-1}) \alpha_n(\mathbf{w}_{0:n})
\end{aligned}$$

where we define $\alpha_n(\mathbf{w}_{0:n}) = \frac{\gamma_n(\mathbf{w}_{0:n})}{\gamma_{n-1}(\mathbf{w}_{0:n-1}) q_n(\mathbf{w}_n|\mathbf{w}_{n-1})}$.
